# Functional characterization of genes mediating cell wall metabolism and responses to plant cell wall integrity impairment

**DOI:** 10.1101/552893

**Authors:** Timo Engelsdorf, Lars Kjaer, Nora Gigli-Bisceglia, Lauri Vaahtera, Stefan Bauer, Eva Miedes, Alexandra Wormit, Lucinda James, Issariya Chairam, Antonio Molina, Thorsten Hamann

**Affiliations:** Institute for Biology, Faculty of Natural Sciences, Norwegian University of Science and Technology, 5 Høgskoleringen, Trondheim, 7491, Norway.; Division of Plant Physiology, Department of Biology, Philipps University of Marburg, 35043 Marburg, Germany.; Division of Cell and Molecular Biology, Department of Life Sciences, Imperial College London, Sir Alexander Fleming Building, South Kensington Campus, London, SW72AZ, UK.; Sjælland erhvervsakademi, Breddahlsgade 1b, 4200 Slagelse, Zealand, Denmark; Laboratory of Plant Physiology, 6708PB Wageningen University and Research, Wageningen, the Netherlands.; Energy Biosciences Institute, University of California, 120A Energy Biosciences Building, 2151 Berkeley Way, MC 5230, Berkeley, CA 94720-5230; Zymergen, Inc. 5980 Horton St, Suite 105 Emeryville, CA 94608, USA; Centro de Biotecnología y Genómica de Plantas, Universidad Politécnica de Madrid (UPM)- Instituto Nacional de Investigación y Tecnología Agraria y Alimentaria (INIA), Campus de Montegancedo-UPM, 28223 Pozuelo de Alarcón, Madrid, Spain.; Departamento de Biotecnología-Biología Vegetal, Escuela Técnica Superior de Ingeniería Agronómica, Alimentaria y de Biosistemas, Universidad Politécnica de Madrid (UPM), 28040 Madrid, Spain.; RWTH Aachen, Institute for Biology I, Worringerweg 3, D-52056 Aachen, Germany.; ADAS, Battlegate Road, Boxworth, Cambridge, CB23 4NN, UK.; Department of Nuclear Safety and Security, International Atomic Energy Agency, Vienna International Centre, PO Box 100, 1400 Vienna, Austria.

**Keywords:** Cell wall, cell wall integrity, cell wall metabolism, cell wall signalling, plant pathogen-interaction

## Abstract

Plant cell walls participate in all plant-environment interactions. Maintaining cell wall integrity (CWI) during these interactions is essential. This realization led to increased interest in CWI and resulted in knowledge regarding early perception and signalling mechanisms active during CWI maintenance. By contrast, knowledge regarding processes mediating changes in cell wall metabolism upon CWI impairment is very limited. To identify genes involved and to investigate their contributions to the processes we selected 23 genes with altered expression in response to CWI impairment and characterized the impact of T-DNA insertions in these genes on cell wall composition using Fourier-Transform Infrared Spectroscopy (FTIR) in *Arabidopsis thaliana* seedlings. Insertions in 14 genes led to cell wall phenotypes detectable by FTIR. A detailed analysis of four genes found that their altered expression upon CWI impairment is dependent on THE1 activity, a key component of CWI maintenance. Phenotypic characterizations of insertion lines suggest that the four genes are required for particular aspects of CWI maintenance, cell wall composition or resistance to *Plectosphaerella cucumerina* infection in adult plants. Taken together, the results implicate the genes in responses to CWI impairment, cell wall metabolism and/or pathogen defence, thus identifying new molecular components and processes relevant for CWI maintenance.

## Introduction

Plant cell walls are involved in all interactions between plants and their environment. Examples include pathogen infection or exposure to drought, where wall composition and structure change to prevent water loss, pathogen susceptibility or at least limit further pathogen spread (Bacete et al., 2018; Novaković et al., 2018). These changes of the walls are exemplified by reinforcement with callose during infection or modifications of pectic polysaccharides to prevent water loss during exposure to drought stress (Chowdhury et al., 2016; Dinakar & Bartels, 2013). Cell walls are extremely plastic, undergoing dynamic changes to enable plant cells to expand and differentiate during growth and development (Bidhendi & Geitmann, 2015). Controlled deposition of cellulose microfibrils through interactions between cellulose synthases and microtubules during cell expansion exemplify the changes in cell wall organization, permitting tightly controlled cell expansion (Gutierrez et al., 2009; Paredez et al., 2006). Deposition of suberin and lignin during formation of the casparian strip in pericycle cells of the primary root exemplifies modifications of cell walls during cell differentiation (Barbosa et al., 2019; Doblas et al., 2017; Lee et al., 2013). These examples illustrate processes active during plant-environment interactions and development, enabling cell walls to fulfill their respective biological functions.

How do cell walls perform these various functions, which sometimes involve opposite performance requirements, while simultaneously maintaining their functional integrity? The available evidence supports the existence of a dedicated mechanism, which is monitoring the functional integrity of the plant cell wall and initiates adaptive changes in cellular and cell wall metabolism to maintain cell wall integrity (CWI) (De Lorenzo et al., 2018; Doblas et al., 2018; Hamann, 2015; Kieber & Polko, 2019; Wolf, 2017). Studies of the mode of action of the CWI maintenance mechanism often investigate the responses to cell wall damage (CWD), which can be generated by cell wall degrading enzymes (cellulase, pectinase etc.) or compounds like isoxaben (ISX) (Engelsdorf et al., 2018). ISX inhibits specifically cellulose production during primary cell wall formation in elongating plant cells (Heim et al., 1990; Scheible et al., 2001; Tateno, Brabham, & DeBolt, 2016). Established responses to CWD include growth inhibition involving cell cycle arrest, changes in the levels of phytohormones like jasmonic acid (JA), salicylic acid (SA) and cytokinins (CKs) as well as changes in cell wall composition involving pectic polysaccharides, lignin and callose deposition (Cano-Delgado et al., 2003; Denness et al., 2011; Ellis & Turner, 2001; Gigli-Bisceglia et al., 2018; Manfield et al., 2004).

The available evidence implicates receptor-like kinases (RLK) like MALE DISCOVERER 1-INTERACTING RECEPTOR LIKE KINASE 2 (MIK2), FEI1, FEI2, THESEUS 1 (THE1) and FERONIA (FER) in CWI maintenance (Engelsdorf et al., 2018; Feng et al., 2018; Hematy et al., 2007; Van der Does et al., 2017; Xu et al., 2008). THE1 and FER belong to the *Catharanthus roseus* RLK1-like kinase (*Cr*RLK1L) family, which has 17 members. These RLKs consist of an intracellular Serine / Threonine-kinase domain, a transmembrane domain and an extracellular domain exhibiting similarity to the malectin domain originally identified in *Xenopus laevis* (Franck et al., 2018). Currently it is not clear if malectin domains in *Cr*RLK1Ls are either required for binding to cell wall epitopes, mediate protein-protein interaction or actually do both (Du et al., 2018; Feng et al., 2018; Gonneau et al., 2018; Haruta et al., 2018; Moussu et al., 2018; Stegmann et al., 2017). FER is required during gametophytic and root hair development, salt stress, JA signaling and coordination between abscisic acid-(ABA) and JA-based signaling processes (Duan et al., 2010; Escobar-Restrepo et al., 2007; Feng et al., 2018; Guo et al., 2018; Kanaoka & Torii, 2010; Shih et al., 2014; Yu et al., 2012; Zhao et al., 2018). MIK2 and THE1 are required for root development, CWD-induced lignin and phytohormone production as well as resistance to the root pathogen *Fusarium oxysporum* (Engelsdorf et al., 2018; Gonneau et al., 2018; Hematy et al., 2007; Van der Does et al., 2017). FEI1 and FEI2 have been originally identified through their impact on seedling root growth on medium containing 4.5% sucrose and subsequently implicated in a cell wall signaling pathway involving the SALT OVERLY SENSITIVE5 (SOS5) and FEI2 (Harpaz-Saad et al., 2011; Shi et al., 2003; Xue et al., 2017). In parallel, ion-channels, like MID1-COMPLEMENTING ACTIVITY 1 (MCA1) and MECHANOSENSITIVE CHANNEL OF SMALL CONDUCTANCE-LIKE 2 (MSL2) and 3 (MSL3) were shown to contribute to activation of CWD-induced responses in plants (Denness et al., 2011; Engelsdorf et al., 2018). MCA1 was originally identified through its ability to partially complement a MID1/CCH1-deficient *Saccharomyces cerevisiae* strain (Nakagawa et al., 2007). In yeast MID1/CCH1 form a plasmamembrane-localized stretch-activated calcium channel required both for mechano-perception and CWI maintenance (Levin, 2011). CWD-induced responses in plants (like in yeast cells) seem also to be sensitive to turgor manipulation (Hamann, 2015; Levin, 2011). The reason being that in *Arabidopsis thaliana* seedlings, exposed simultaneously to ISX and mild hyperosmotic conditions, most of the CWD-induced responses are suppressed in a concentration dependent manner (Engelsdorf et al., 2018; Hamann et al., 2009). The early signals generated seem to be conveyed to downstream response mediators through changes in production of reactive oxygen species (ROS) and phytohormones (JA/SA/CKs) (Denness et al., 2011; Gigli-Bisceglia et al., 2018). Enzymes implicated in ROS production upon CWI impairment are NADPH-oxidases like RESPIRATORY BURST OXIDASE HOMOLOGUE (RBOH) D/F (after ISX-treatment) or RBOH H/J during pollen tube development (Jiménez-Quesada et al., 2016). NADPH-oxidase activity in turn can be regulated via calcium binding, differential phosphorylation involving kinases controlled by changes in calcium levels (CALCINEURIN INTERACTING KINASE 26, CIPK26), activated in response to pathogen infection through phosphorylation involving BOTRYTIS INDUCED KINASE 1 (BIK1) or controlled via RHO GTPases, a ROPGEF and FER (Duan et al., 2010; Han et al., 2018; Kadota et al., 2014).

This abbreviated overview of molecular components active during plant CWI maintenance illustrates the increase in knowledge regarding putative CWI sensors and early signal transduction elements in recent years. Whilst it is fascinating to know about early CWD perception and signaling processes we also need to understand how signals generated lead to changes in cell wall composition and structure to dissect the mode of action of the CWI maintenance mechanism thoroughly. This is of particular interest in the context of targeted modification of biomass quality and improvement of food crop performance since the CWI maintenance mechanism seems to be an important component of cell wall plasticity (Doblin et al., 2014; Mahon & Mansfield, 2019). Cell wall plasticity in turn has been discussed as the root cause for the apparently limited success of efforts aimed at optimizing biomass quality that have been achieved so far (Doblin et al., 2014).

We wanted to identify additional components and molecular processes, which are mediating responses to CWD and adaptive changes in cell wall metabolism. To achieve these aims we selected candidate genes using microarray-based expression profiling data deriving from ISX-treated Arabidopsis seedlings. Fourier Transform Infrared (FTIR) Spectroscopy was then used to identify candidate genes where insertions lead to cell wall changes on the seedling level. We performed in depth studies for four genes to validate the approach. These studies involved confirming that gene expression is responsive to ISX, determining if expression is controlled by THE1 and investigating how loss of function alleles for these genes affect cell wall composition in adult plants, resistance to the necrotrophic pathogen *Plectosphaerella cucumerina* and responses to ISX-induced CWD impairing CWI.

## Materials and Methods

### Reagents

All chemicals and enzymes were purchased from Sigma-Aldrich unless stated otherwise.

### Plant Material

Wild-type and mutant *Arabidopsis thaliana* lines used in this study were ordered from the Nottingham Arabidopsis Stock Centre (http://arabidopsis.info/). Detailed information is listed in Supplemental Table S1. Seedlings were grown for 6 days in liquid culture (2.1 g/L Murashige and Skoog Basal Medium, 0.5 g/L MES salt and 1 % sucrose at pH 5.7) before treatment with 600 nM isoxaben (in DMSO) as described (Engelsdorf et al., 2018). For cell wall analysis, plants were grown on soil (Pro-Mix HP) in long-day conditions (16 h light, 11000 Lux, 22°C; 8 h dark, 20 °C; 70 % relative humidity). For pathogen infection assays, plants were grown in phytochambers on sterile soil-vermiculite (3:1) under short-day conditions (10 h of light/14 h of dark) at 20-21 °C.

### Pathogen Infection Assays

For *Plectosphaerella cucumerina* BMM (*Pc*BMM) pathogenicity assays, 18 days-old plants (n >15) were sprayed with a spore suspension (4 x 10^6^ spores/ml) of the fungus as previously described (Delgado-Cerezo et al., 2011; Sanchez-Vallet et al., 2010). Fungal biomass *in planta* was quantified by determining the level of the *Pc*BMM β-tubulin gene by qPCR (forward primer: CAAGTATGTTCCCCGAGCCGT and reverse primer: GGTCCCTTCGGTCAGCTCTTC) and normalizing these values to those of *UBIQUITIN-CONJUGATING ENZYME21* (*UBC21, AT5G25760*).

### Quantitative RT-PCR

Total RNA was isolated using a Spectrum Plant Total RNA Kit (Sigma-Aldrich). Two micrograms of total RNA were treated with RQ1 RNase-Free DNase (Promega) and processed with the ImProm-II Reverse Transcription System (Promega) for cDNA synthesis. qRT-PCR was performed with a Roche LightCycler 480 system using LightCycler 480 SYBR Green I Master. Gene expression levels were determined as described (Gigli-Bisceglia et al., 2018). The following gene-specific primers have been used for time course expression analysis in Col-0:

*ACT2-FOR* (5’-CTTGCACCAAGCAGCATGAA-3’),

*ACT2-REV* (5’-CCGATCCAGACACTGTACTTCCTT-3’),

*WSR1-FOR* (5’-TATGGTGATGAACTTTGCGTTC-3’),

*WSR1-REV* (5’-ACTCAACAGTAGCATCTCCTGA-3’),

*WSR1A-FOR* (5’-TACGCTGCTACTGGTCAACG-3’),

*WSR1A-REV* (5’-TTCCTCCAATCACCGGCATC-3’),

*WSR2-FOR* (5’-CTCACTTCCATCGTTTCAAGTG-3’),

*WSR2-REV* (5’-GAAACCAAACGTGGCCTAAA-3’),

*WSR3-FOR* (5’-GAAAGCACGAGACTGGAACG-3’),

*WSR3-REV* (5’-TATCCACCCTCCAACGCAAA-3’),

*WSR4-FOR* (5’-AGCCCTGAGAGATCAAGCATT-3’),

*WSR4-REV* (5’-AGCTCAACTAAGCGATGAAGC-3’).

For the characterization of T-DNA insertion lines, the following primers have been employed: *ACT2-FOR, ACT2-REV, WSR1-FOR, WSR1-REV, WSR2-FOR, WSR2-REV, WSR3-FOR2* (5’-TCTTATCCGGTTGCGGAAGG-3’),

*WSR3-REV2* (5’-GTGGTGAGATGACCCAGAGC-3’),

*WSR4-FOR2* (5’-CTTGATGCAGTTGTGAAAGCA-3’),

*WSR4-REV2* (5’-TCTTCACCGAAACAATCATCC-3’).

### FTIR Spectroscopy and Analysis

For FTIR analysis, 4 biological replicates per genotype and 5 technical replicates per biological replicate were collected (i.e. for each genotype 20 spectra were collected). Spectra for each technical replicate were measured from 800 to 5000 cm^-1^ with 15 accumulations per measurement on a Bruker Vertex 70. All spectra were measured at 10 kHz, with a 10 kHz lowpass filter and the Fourier transform was carried out using Blackman-Harris 3-term. Atmospheric compensation was carried out on the data using OPUS version 5 (www.bruker.com). The spectra were cropped to the area between 802 cm^-1^ to 1820 cm^-1^ to cover informative wavenumbers as described in (Mouille et al., 2003). Linear regression was carried out based on the first ten points in either end of the spectra and used for baseline correction. The data was normalized to sum 1 with any negative values still present set to 0 for normalization purposes. Biological variation in the Col-0 controls was determined based on three independent experiments carried out with 4 biological replicates and 5 technical replicates per biological replicate (i.e. 20 spectra per experiment). The difference between the insertion lines and Col-0 was calculated by averaging all the technical repeats for a line and subtracting the corresponding average from Col-0. The difference between the insertion lines and Col-0 was plotted by wavelength. Two times the standard deviation of Col-0 was chosen as a cutoff as it would indicate significance if the natural variation is assumed to be symmetrical across Col-0 and the insertion line.

### Cell Wall Analysis

Cell wall preparation and analysis were performed as described (Yeats et al., 2016) with minor modifications. For analysis of stem cell wall composition, major stems of three plants per genotype were pooled to form one biological replicate. For analysis of leaf cell wall composition, whole leaf rosettes of three plants per genotype were pooled to form one biological replicate. Four biological replicates were analysed in all cases. Plant samples were immediately flash-frozen in liquid nitrogen after sampling and lyophilized. Dried material was ball-milled with zirconia beads in a Labman robot (www.labmanautomation.com), extracted three times with 70 % ethanol at 70°C and dried under vacuum. Starch was removed using a Megazyme Total Starch Kit according to the manufacturer’s instructions. After drying under vacuum, de-starched alcohol insoluble residue (AIR) was weighed out in 2 ml screw caps tubes for cell wall monosaccharide analysis (2 mg AIR) and GC vials for lignin analysis (1.2 mg AIR), respectively, with the Labman robot (0.2 mg tolerance). Cellulose, neutral sugars and uronic acids were determined following the published one-step two-step hydrolysis protocol (Yeats et al., 2016). High-performance anion-exchange chromatography with pulsed amperometric detection (HPAEC-PAD) was performed on a Thermo Fisher Dionex ICS-3000 system with CarboPac PA-20 and PA-200 columns as described (Yeats et al., 2016). Acetyl bromide soluble lignin was quantified as described (Chang et al., 2008)

### Phytohormone Analysis

JA and SA were extracted and analysed as described (Engelsdorf et al., 2018). Briefly, extraction was performed in 10 % methanol / 1 % acetic acid with Jasmonic-d_5_ Acid and Salicylic-d_4_ Acid (CDN Isotopes) as internal standards. Quantification was performed on a Shimadzu UFLC XR / AB SCIEX Triple Quad 5500 system using the following mass transitions: JA 209 > 59, D_5_-JA 214 > 62, SA 137 > 93, D_4_-SA 141 > 97.

### Lignin Detection in Roots

Lignification in seedling roots (n>15) was analysed 24 h after start of treatment. Lignified regions were detected with phloroglucinol-HCL, photographed with a Zeiss Axio Zoom.V16 stereomicroscope and quantified as described (Engelsdorf et al., 2018).

### Statistical Analysis

Statistical significance was assessed using Student’s *t*-test in Microsoft Excel (2-tailed distribution, two-sample unequal variance). Statistically significant differences are indicated by * *p* < 0.05, * *p* < 0.01. Boxplots were generated using R package “boxplot” with default settings (range = 1.5*IQR).

## Results

### Identification of candidate genes

Previously, we have performed time course experiments to characterize the response of Arabidopsis seedlings to ISX-induced CWD (Hamann et al., 2009). Affymetrix ATH1 microarrays were used to detect changes in transcript levels up to 36 hours after start of ISX treatment. The phenotypic characterization of seedlings detected lignin deposition in root tips and enhanced JA production after 4-6 hours of ISX-treatment (Hamann et al., 2009). Based on these results we hypothesized that genes exhibiting transcriptional changes after 4 hours might be involved in CWD responses (phytohormone and lignin production) as well as cell wall modifying processes in general. Analysis of the microarray derived expression data suggested that the transcript levels of several hundred genes change after 4 hours of exposure to ISX. We used public expression data (www.genevestigator.com) to identify genes exhibiting differential expression after 4 hours of ISX treatment and elevated expression in tissue types where cell wall modification or production occurs preferentially (primary root elongation zone and expanding hypocotyl). To determine whether these candidates are involved in cell wall related processes, we decided to perform a pilot study characterizing the phenotypes of T-DNA insertion lines for 23 candidate genes. Supporting information Table 1 (Table S1) lists the 23 candidate genes with their database annotations, the probe sets representing the genes on the Affymetrix ATH1 microarray and insertion lines used for characterization. Figure S1 summarizes the microarray-derived expression data for the 23 candidate genes generated in the original time course expression profiling experiments. The genes are separated based on their transcript levels either being apparently increased (Figure S1a) or decreased (Figure S1b) over time. Figure S2 illustrates the putative transcript levels of the candidate genes in different tissues / organs. The genevestigator-derived expression data suggest that several candidate genes are involved in cellular and biological processes, which affect or involve plant cell wall metabolism.

**Table 1:** Candidate genes selected from the transcriptomics / FTIR-based screen. Gene annotations are based on Araport11 and references listed. WSR: Wall Stress Response.

|  | AGI | Gene Annotation | Reference |
| --- | --- | --- | --- |
| **WSR1** | *At3g13650* | *DIR7*, Disease resistance-responsive (dirigent-like protein) family protein | Paniagua et al., 2017 |
| **WSR2** | *At2g35730* | Heavy metal transport / detoxification superfamily protein | De Abreu-Neto et al., 2013 |
| **WSR3** | *At5g47730* | *SFH19*, Sec14p-like phosphatidylinositol transfer family protein | De Campos et al., 2017 |
| **WSR4** | *At2g41820* | *PXC3*, Leucine-rich repeat protein kinase family protein | Wang et al., 2013 |

### FTIR-based analysis detects cell wall phenotypes in mutant seedlings

Performing detailed cell wall analysis for insertion lines in 23 candidate genes would be time consuming and possibly not very efficient. Previously, FTIR has been successfully used as an efficient approach to classify *Arabidopsis* mutants with altered cell wall architecture (Mouille et al., 2003). We used this approach as foundation to facilitate identification of insertions in candidate genes leading to changes in cell wall composition or structure in Arabidopsis seedlings. FTIR spectra were collected for analysis from total cell wall material derived from 6 days-old, liquid culture grown Col-0 seedlings or seedlings with T-DNA insertions in the candidate genes. Initially only Col-0 samples were characterized to establish the variability observed in controls. Subsequently, twice the standard deviation of the Col-0 variability was used as a cut-off to identify insertions in candidate genes causing significant changes in the FTIR spectra. Based on this criterium FTIR spectra for 14 of the 23 insertion lines analyzed exhibited significant differences (Figure S3). Pronounced differences were observed for insertions in *At5g24140* (*SQUALENE MONOOXYGENASE2, SQP2)* and *At5g49360* (*BETA-XYLOSIDASE1, ATBXL1*) in the 1700-1600 cm^-1^ (pectin ester, carboxylate / carbonyl side groups) and 1200-950 cm^-1^ (characteristic for cellulose and elements of pectic polysaccharides) areas (Figure S3, blue rectangle) (Arsovski et al., 2009; Rasbery et al., 2007; Szymanska-Chargot et al.,. Insertion lines for *At2g41820* (*PHLOEM INTERCALATED WITH XYLEM / TRACHEARY ELEMENT DIFFERENTIATION INHIBITORY FACTOR RECEPTOR-CORRELATED 3, PXC3*), *At3g11340 (UDP GLYCOSYLTRANSFERASE 76B1, UGT76B1*), *At4g33420* (*PEROXIDASE 47, PRX47*), *At4g35630* (*PHOSPHOSERINE AMINO-TRANSFERASE 1, PSAT 1*), *At5g48460* (*FIMBRIN 2, ATFIM2*), *At5g47730* (*SEC14-HOMOLOGUE 19, SFH19*) and *At5g65390* (*ARABINOGALACTAN PROTEIN 7, AGP7*) exhibited differences in the 1367-1200 cm^-1^ area typical for certain cellulose elements, hemicelluloses and pectins (Figure S3, red rectangles) (Maksym et al., 2018; Seifert, 2018; Szymanska-Chargot et al., 2015; Tokunaga et al., 2009; Wang et al., 2013; Wulfert & Krueger, 2018; Zhang et al., 2016). In the 1200-950 cm^-1^ area distinctive differences were detected for insertion lines in five candidate genes *At1g07260* (*UGT71C3*), *At1g74440, At2g35730, At3g13650 (DIRIGENT PROTEIN 7, DIR7)* and *At4g33300* (*ACTIVATED DISEASE RESISTANCE-LIKE 1, ADR1-L1*) (Figure S3, purple rectangles) (Dong et al., 2016; Meier et al., 2008; Paniagua et al., 2017; Rehman et al., 2018; Wuest et al., 2010). The results from the FTIR-based analysis of the insertion lines suggested that cell wall composition or structure is affected in seedlings with insertions for 14 of the 23 candidate genes examined. The insertions seemed to have distinct effects on cell wall composition / structure based on their apparent separation into three groups.

### Four candidate genes are selected for more detailed characterization

We selected four candidate genes for a more detailed analysis (*At3g13650, At2g35730, At5g47730* and *At2g41820*, Table 1) because of the limited knowledge available regarding their biological functions and the four insertions leading to two qualitatively different FTIR phenotypes. This enabled us to determine also if insertion lines resulting in similar FTIR cell wall phenotypes on the seedling level exhibit similar cell wall phenotypes on the adult plant level. While *At3g13650* and *At2g35730* FTIR-spectra (orange, yellow) seemed to deviate from Col-0 controls mainly in areas characteristic for cellulose and certain types of pectins, *At5g47730* and *At2g41820* spectra (green, blue) deviated mainly in areas characteristic for cellulose elements, hemicelluloses and pectins (Figure 1, numbers highlight wavenumbers diagnostic for certain bands in the spectra according to Szymanska-Chargot et al., 2015). These four genes were classified as WALL STRESS RESPONSE genes or WSRs. *At3g13650* (*WSR1, DIR7*) belongs to a family of disease resistance responsive proteins, which have been implicated in lignan biosynthesis and formation of the casparian strip (Barbosa et al., 2019; Paniagua et al., 2017). Analysis of the available data suggests that *At2g35730* (*WSR2*) is expressed in the female gametophyte and encodes a heavy metal transport / detoxification superfamily protein (De Abreu-Neto et al., 2013; Wuest et al., 2010). Protein homology suggests that *At5g47730* (*WSR3, SFH19*) is related to SEC14 proteins from *S. cerevisiae*, which have been implicated in polarized vesicle transport (KF de Campos & Schaaf, 2017). *At2g41820* (*WSR4, PXC3*) encodes a putative leucine-rich repeat receptor kinase, belonging to a family where other members have been implicated in organization of secondary vascular tissue (Wang et al., 2013).

**Figure 1.**
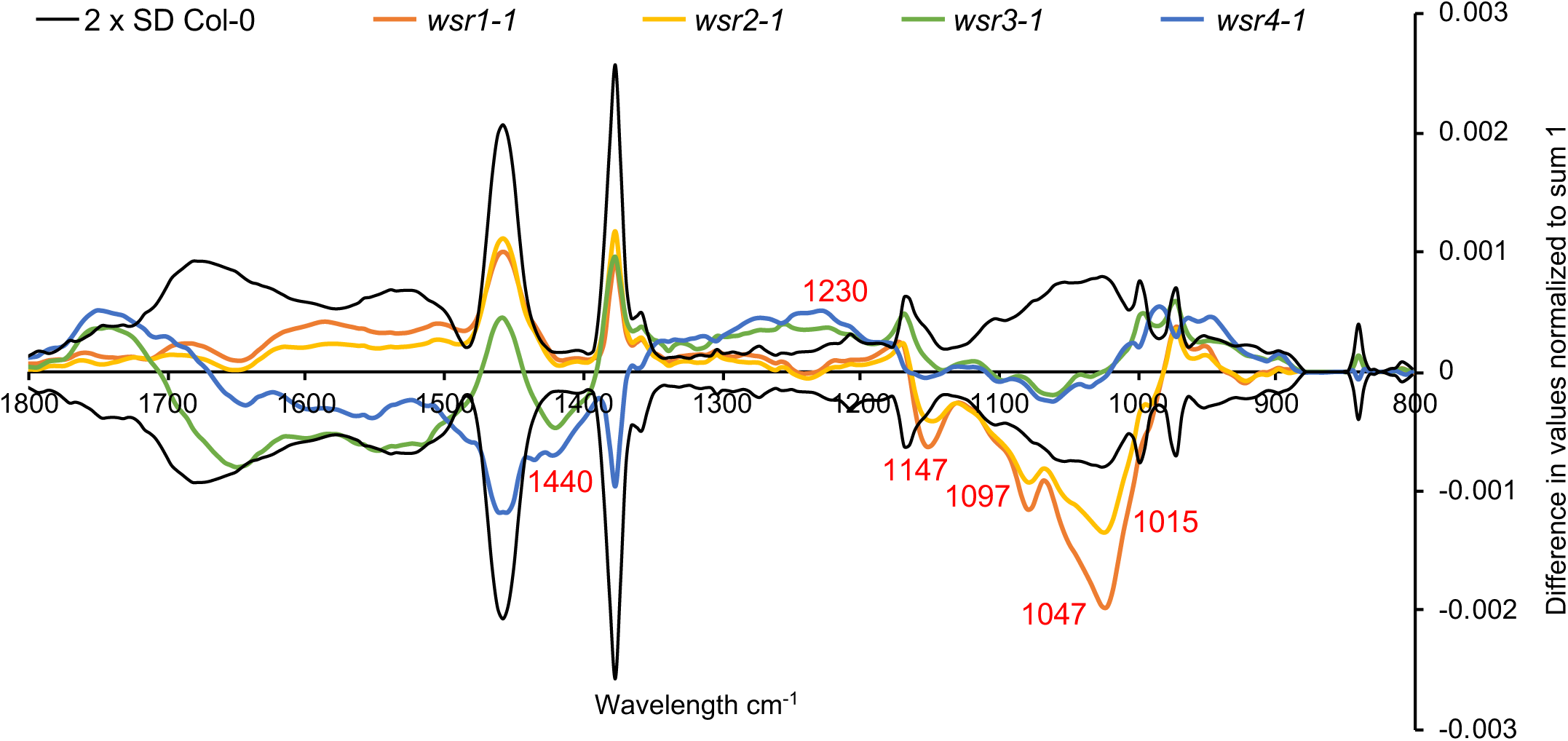
Overview of average Fourier-Transform Infrared (FTIR) spectra from Col-0, *wsr1-1, wsr2-1, wsr3-1* and *wsr4-1* seedlings. Black lines indicate 2 x SD for Col-0-derived material. The differently coloured lines for *wsr1-1* (*DIR7*), *wsr2-1* (*At3g35730*), *wsr3-1* (*SFH19*) and *wsr4-1* (*PXC3*) are based on the normalized average FTIR spectra for Col-0 seedlings minus the average spectra of the individual mutant. Numbers in red indicate bands in the infrared spectra of plant cell wall material indicative for certain classes of cell wall polysaccharides.

### Quantitative gene expression analysis confirms transcriptomics results and suggests THE1 is controlling *WSR* gene expression

DNA microarray-based expression analysis of Arabidopsis seedlings suggested that the transcript levels of the *WSR* genes are changing in response to ISX treatment (Figure S1). To confirm this, we performed time course experiments and transcript levels of the four genes were determined through quantitative reverse transcription – polymerase chain reaction (qRT - PCR) in mock- or ISX-treated seedlings. The transcript levels of *WSR1, 2* and *3* increased after 4 hours of ISX treatment and remained elevated compared to mock controls (Figure 2a). Transcript levels of *WSR4* were reduced after an initial transient increase. To establish if expression of the *WSR* genes is controlled by the THE1-mediated CWI maintenance mechanism we investigated *WSR* transcript levels in THE1 loss (*the1-1*) - or gain-of-function (*the1-4*) seedlings (Hematy et al., 2007; Merz et al., 2017). In ISX-treated Col-0 seedlings *WSR1, 2* and *3* transcript levels were increased while *WSR4* seemed slightly reduced after eight hours (Figure 2b). *WSR1* transcript levels were not increased in *the1-1* seedlings but the increase was enhanced in *the1-4*. *WSR2* transcript levels changed in *the1-1* seedlings as in Col-0 while the increase was enhanced in *the1-4*. Increases in *WSR3* expression were apparently slightly reduced in *the1-1* seedlings compared to Col-0 while the increase was again more pronounced in *the1-4* than in ISX-treated Col-0 seedlings. Decrease of *WSR4* expression seemed absent in *the1-1* compared to ISX-treated Col-0 seedlings and was enhanced in ISX-treated *the1-4* seedlings. In conclusion, both DNA microarray and qRT-PCR expression analyses showed that transcript levels of *WSR1, 2* and *3* increased while *WSR4* decreased in ISX-treated seedlings over time. *WSR* transcript levels seem to be influenced to different degrees by changes in THE1 activity with increased THE1 activity affecting all WSR genes while decreased THE1 activity seems to affect particularly strongly *WSR1*.

**Figure 2.**
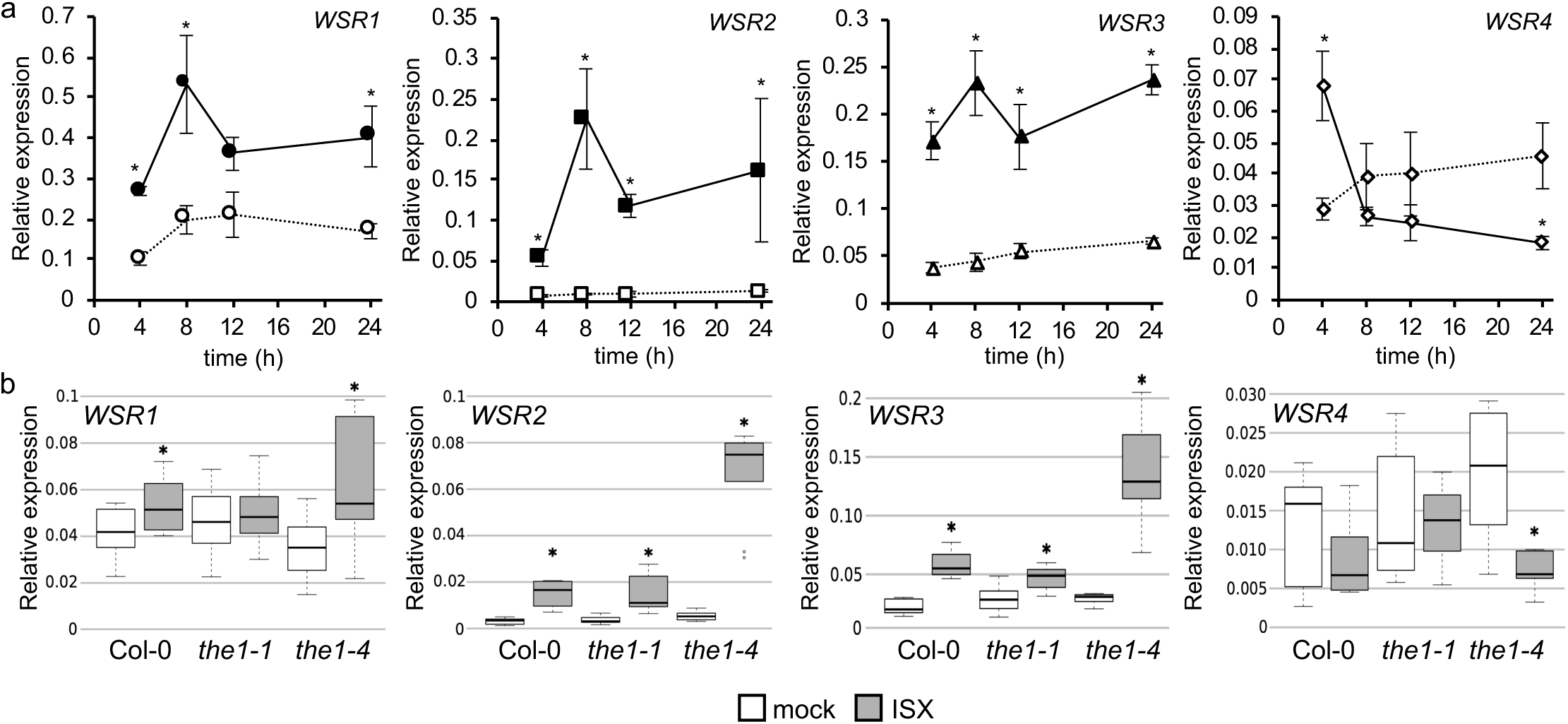
Candidate gene expression profiling in seedlings exposed to cellulose biosynthesis inhibition. **(a)** Gene expression of *WSR1, 2, 3* and *4* in Col-0 seedlings at the indicated time points after mock (empty symbols, dotted lines) or ISX (filled symbols, solid lines) treatment according to qRT-PCR analysis. Values were normalized to *ACT2* and represent means from 3 independent experiments (n=9). Error bars indicate SD. Asterisks indicate statistically significant differences (*p < 0.05) to mock controls according to Student‘s t test. (**b**) Transcript levels of *WSR1, 2, 3* and *4* in Col-0, *the1-1* and *the1-4* seedlings mock (DMSO) or ISX-treated for 8 hours. Values were normalized to *ACT2* and represent means from 3 independent experiments (n= 8-9). Asterisks indicate statistically significant differences to mock controls according to Student‘s t test (*p < 0.05). The boxes in the boxplot indicate interquartile range (IQR, between 25^th^ and 75^th^ percentile) and the black line in the middle of the box marks the median. The whiskers indicate data points furthest from the median, if they are still within 1.5xIQR from the closest quartile. The data points outside this range are plotted individually.

### Identification of knockout and knockdown alleles for *WSR* genes

Using knockout (KO) or knockdown (KD) alleles generated through T-DNA insertions is a well-established and successful method to characterize genes of interest (Alonso et al., 2003). We identified two independent T-DNA insertion lines for each of the four genes using the Arabidopsis Gene Mapping Tool (http://signal.salk.edu/cgi-bin/tdnaexpress). Plants homozygous for the insertions were isolated using PCR-based genotyping as well as insertion positions in the individual gene and their effects on transcript levels determined. For *WSR1* the first insertion is located in the 5’ (*wsr1-1,* Salk_046217) and the second (*wsr1-2,* Salk_092919) in the 3’ untranslated region of the gene (Figure S4a). The insertions in *WSR2* were mapped to the first intron (*wsr2-2,* Salk_123509) and the third exon (*wsr2-1,* Salk_058271) (Figure S4b). For *WSR3* the insertions were located either in the promoter region (*wsr3-2*, SALK_079548) or in the 10^th^ intron (*wsr3-1*, SALK_039575) (Figure S4c). In the *wsr4-2* (SALK_121365) allele the insertion is located in the 1^st^ while the one giving rise to the *wsr4-1* (SALK_082484) allele is located in the 2^nd^ exon (Figure S4d). To determine if the insertions affect transcript levels of the genes, we performed qRT-PCR using total RNA isolated from mock-treated 7 days-old seedlings. These experiments identified four bona fide KO-(*wsr2-1, wsr2-2, wsr3-1, wsr4-2*) and four KD-alleles (*wsr1-1, wsr1-2, wsr3-2* and *wsr4-1*) for the candidate genes (Figure S4a-d).

### Responses to ISX-induced CWD are modified in seedlings with insertions in *WSR1* or *4*

ISX-induced CWD leads to lignin deposition in seedling root tips as well as increased JA and SA production (Cano-Delgado et al., 2003; Ellis & Turner, 2001; Hamann et al., 2009). Here we investigated if the KO / KD alleles isolated for the candidate genes affect these responses by performing experiments with mock or ISX-treated seedlings in liquid culture. After treating the seedlings for seven hours we measured both JA and SA levels (Figure 3a, b). In ISX-treated *wsr4-2* seedlings, we detected reduced production of JA while SA levels were lower in *wsr1-2* seedlings. Lignin was detected after 24 h of ISX treatment using phloroglucinol and quantified using image analysis (Engelsdorf et al., 2018). Mock-treated seedling roots did not show any lignin deposition (Figure 4, Figure S5). We only detected a significant reduction in ISX-treated *wsr4-2* seedlings in lignification after ISX-treatment. These results suggested that *WSR1* and *4* might be involved in the response to ISX-induced CWD.

**Figure 3.**
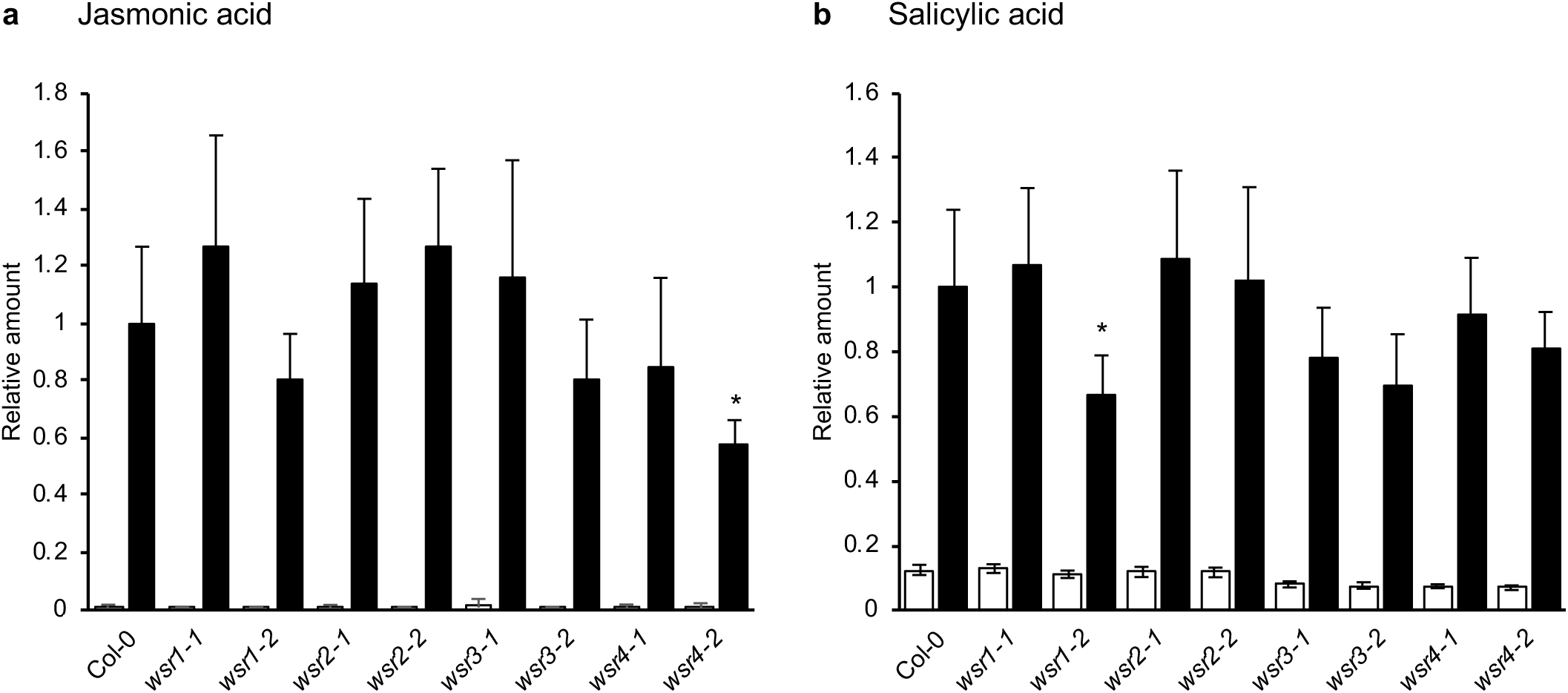
Relative jasmonic acid and salicylic acid accumulation in *wsr* seedlings after ISX treatment. **(a)** Jasmonic acid and **(b)** salicylic acid were quantified in Col-0, *wsr1-1, wsr1-2, wsr2-1, wsr2-2, wsr3-1, wsr3-2, wsr4-1* and *wsr4-2* seedlings after 7 h of mock (empty bars) or ISX (filled bars) treatment. Bars represent mean values from 3-4 independent experiments and error bars indicate SD. Asterisks indicate statistically significant differences to the ISX-treated wild type according to Student‘s t test (*p < 0.05).

**Figure 4.**
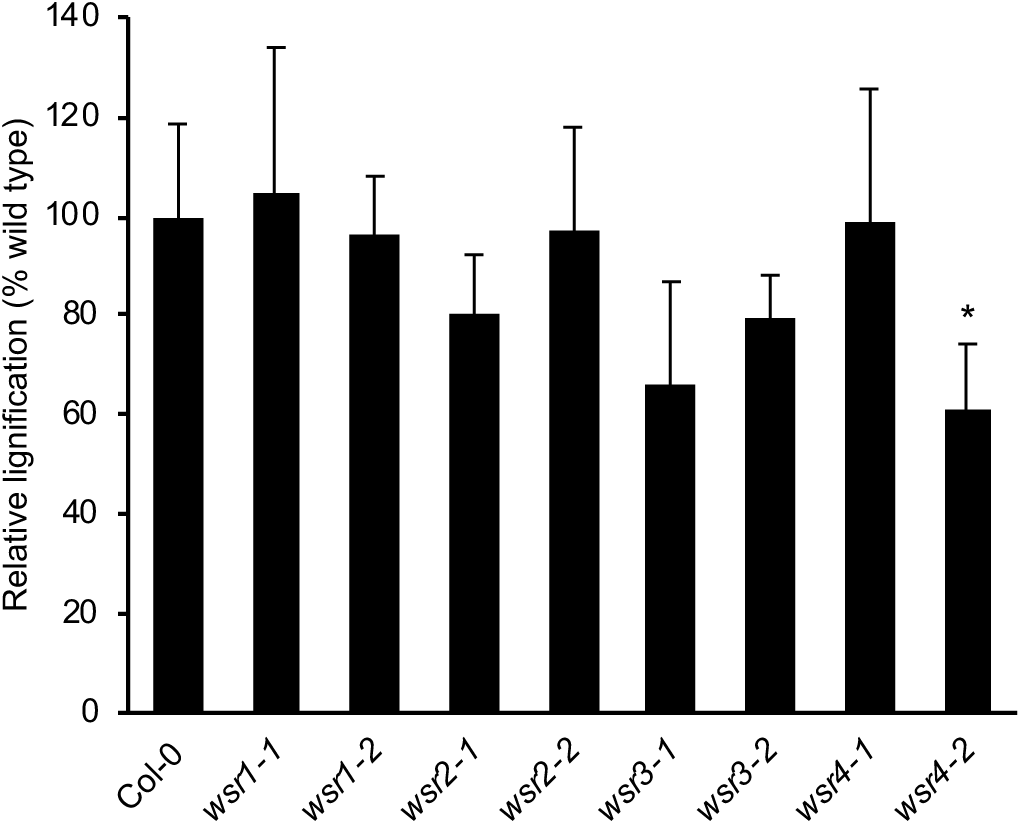
Relative lignification in *wsr* root tips after ISX treatment. Lignification in root tips of Col-0, *wsr1-1, wsr1-2, wsr2-1, wsr2-2, wsr3-1, wsr3-2, wsr4-1* and *wsr4-2* seedlings was quantified after 24 h of ISX treatment. Bars represent mean values from 3 independent experiments and error bars indicate SD. Asterisks indicate statistically significant differences to the wild type according to Student‘s t test (*p < 0.05).

### *WSR1, 2, 3* and *4* contribute to cell wall formation during stem growth

To determine whether the genes of interest affect cell wall metabolism in general, the levels of cellulose, uronic acids and neutral cell wall sugars were determined in cell wall preparations from rosette leaves and mature stems of Col-0 and *wsr* mutant plants. Here we only investigated the strongest KO or KD allele for each candidate gene. We detected a reduction in cellulose content only in leaves of *wsr4-2* plants compared to Col-0 (Figure 5a). Our quantification of cellulose in mature stems detected increased amounts in *wsr1-2* and *wsr2-1*, while cellulose was reduced in *wsr4-2* stems (Figure 5b). Analysis of lignin content in stems of adult plants did not detect any differences between Col-0 and mutant plants (Figure S6). Analysis of neutral cell wall sugars and uronic acids in leaves detected only for the low-abundant glucuronic acid in *wsr1-2* significant differences to Col-0 controls (Figure 6a). In stem-derived material, we observed enhanced levels of rhamnose and xylose in *wsr1-2* (Figure 6b). In *wsr3-1* stems fucose, rhamnose, arabinose and galactose contents were elevated. In *wsr4-2* glucose amounts were reduced while mannose was slightly enhanced compared to Col-0 controls. To summarize, our cell wall analyses detected differences in cellulose and different neutral cell wall sugar contents with effects detected most pronounced in stems. The similarities and variability observed regarding cell wall phenotypes suggest that the different genes may be involved in distinct but also overlapping aspects of cell wall metabolism in adult plants.

**Figure 5.**
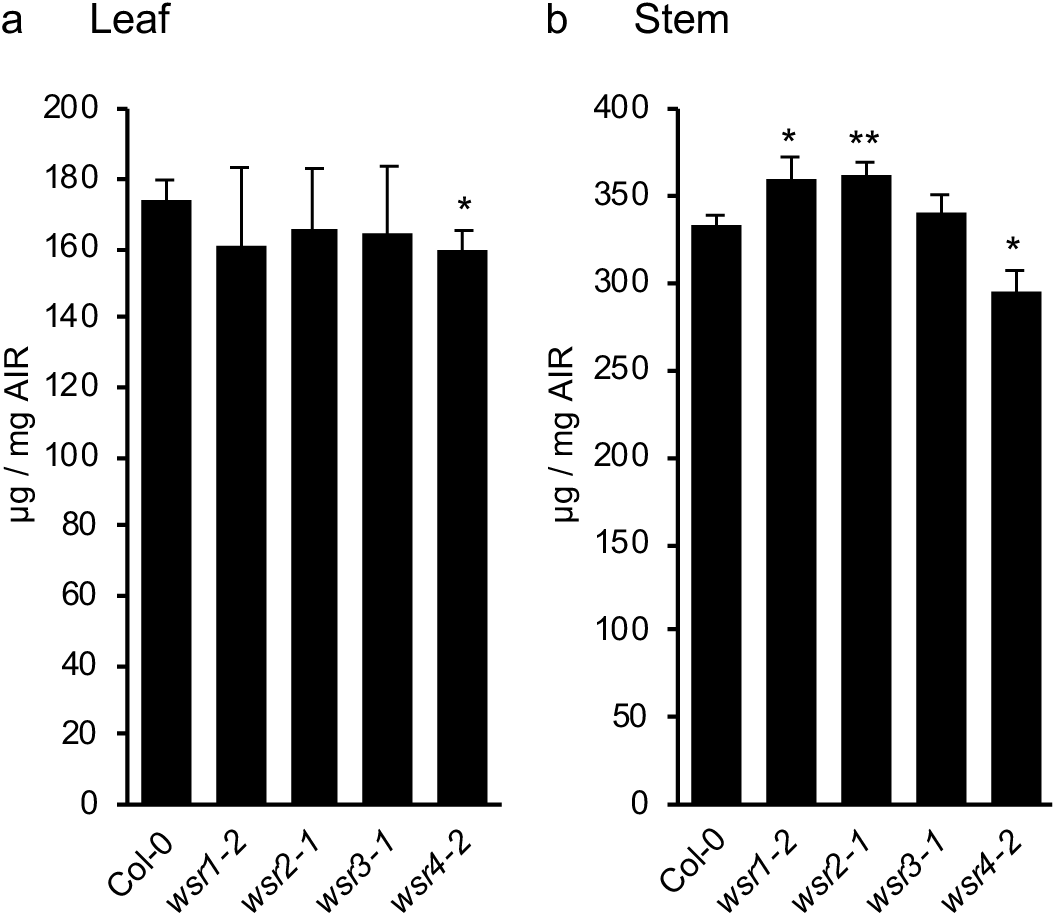
Cellulose content in adult *wsr* plants. Cellulose content was quantified in cell wall preparations from 5 weeks-old Col-0, *wsr1-2, wsr2-1, wsr3-1* and *wsr4-2* plants. **(a)** Leaf cellulose, **(b)** stem cellulose. Bars represent mean values and error bars indicate SD (n = 4). Asterisks indicate statistically significant differences to the wild type according to Student‘s t test (*p < 0.05; **p < 0,01).

**Figure 6.**
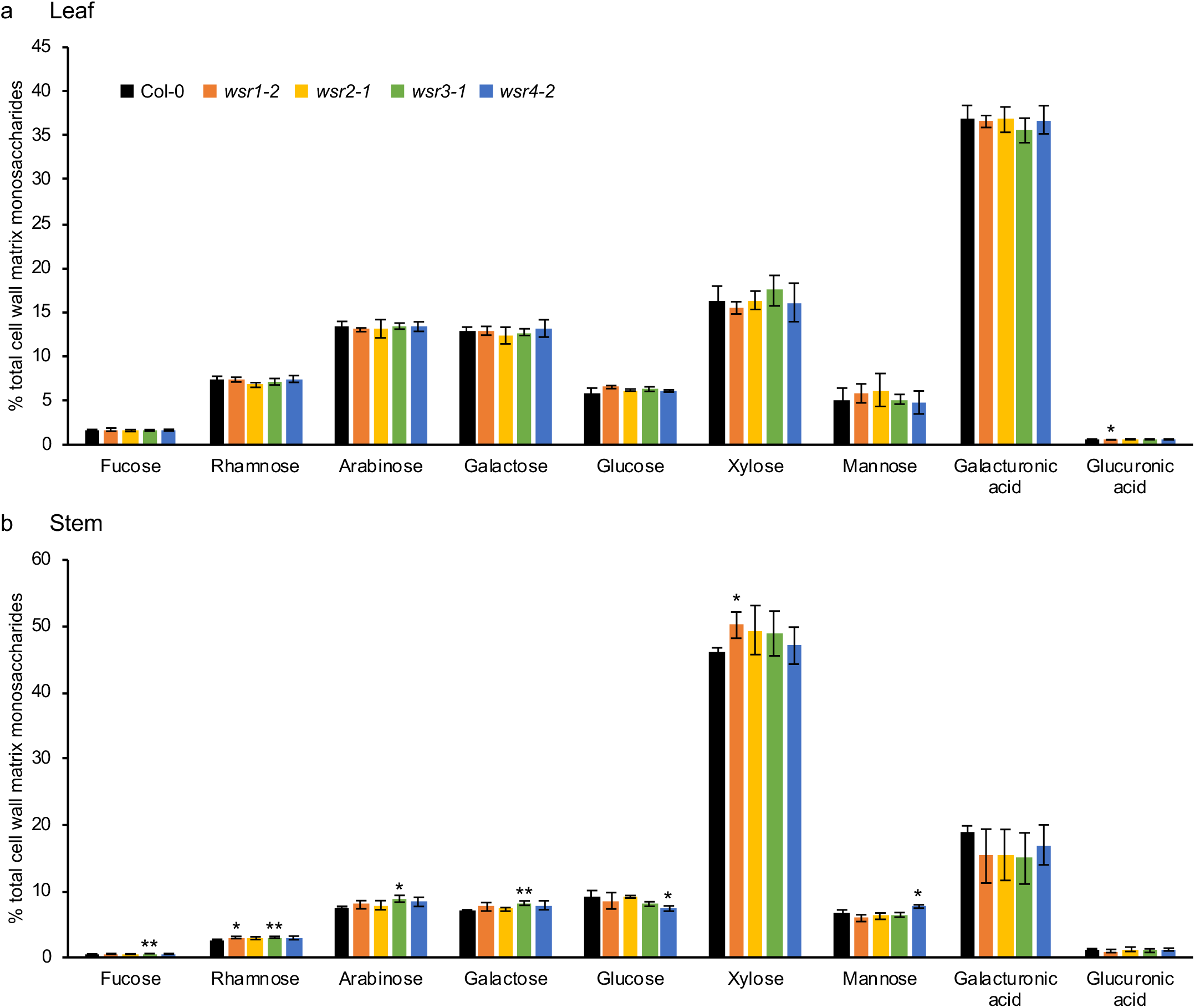
Cell wall matrix monosaccharide composition in adult *wsr* plants. Relative amounts of the monosaccharides Fucose, Rhamnose, Arabinose, Galactose, Glucose, Xylose, Mannose, Galacturonic acid and Glucoronic acid were quantified in cell wall matrix hydrolysates of 5 weeks-old Col-0, *wsr1-2, wsr2-1, wsr3-1* and *wsr4-2* plants. **(a)** Leaf monosaccharides, **(b)** stem monosaccharides. Bars represent mean values and error bars indicate SD (n = 4). Asterisks indicate statistically significant differences to the wild type according to Student‘s t test (*p < 0.05; **p < 0,01).

### Plants with insertions in *WSR1* exhibit pathogen response phenotypes

It has been shown previously that the CWI monitoring RLKs *THE1* and *MIK2* affect pathogen susceptibility (Van der Does et al., 2017). Here we investigated if the mutations in WSR genes also affect the outcome of plant-pathogen interactions by inoculating adult plants carrying insertions in the four genes with the necrotrophic fungus *Plectosphaerella cucumerina* BMM (*Pc*BMM) and quantifying fungal biomass five days post inoculation (Figure 7). *Arabidopsis Gβ 1* (*agb1-1*) and *irregular xylem 1* (*irx1-6*) plants were included as controls since they exhibit reduced (*agb1-1*) or enhanced resistance (*irx1-6*) to *Pc*BMM infection (Hernandez-Blanco et al., 2007; Llorente et al., 2005). Fungal growth on infected *wsr2-1, 3-1* and *4-2* plants was similar to Col-0 controls. However, growth was significantly enhanced on *wsr1-2* plants, suggesting that reduction of *WSR1* gene expression affects resistance to *Pc*BMM infection and implicating this gene in disease resistance response.

**Figure 7.**
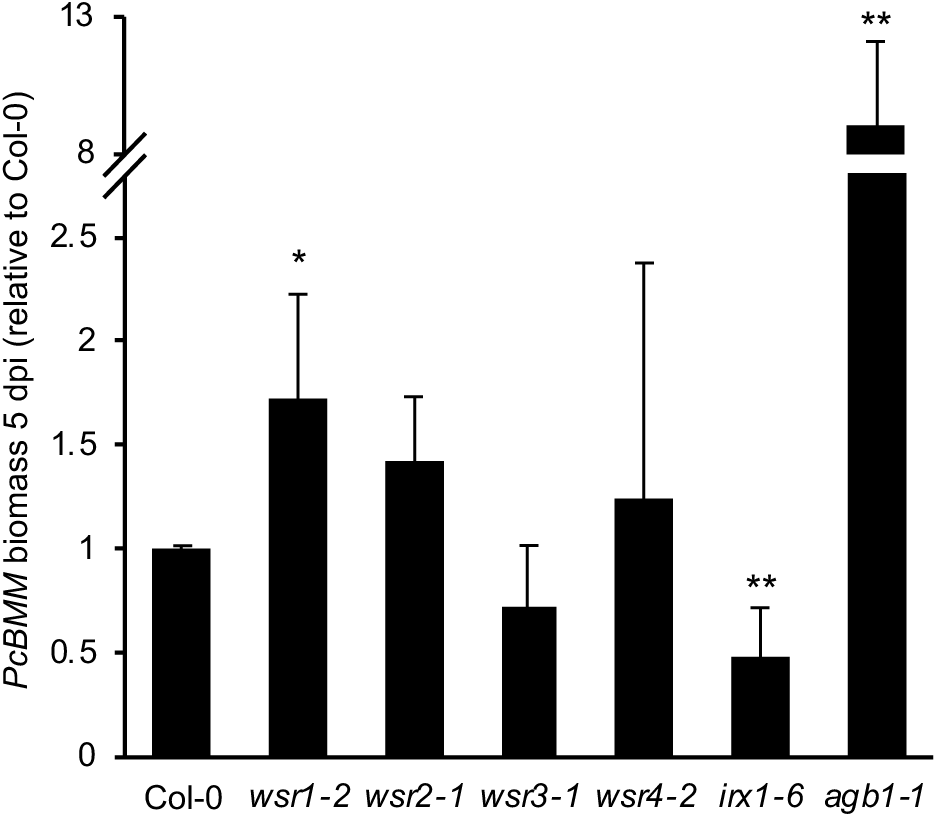
Relative susceptibility of *wsr* plants to *Plectosphaerella cucumerina*. 3 weeks-old Col-0, *wsr1-2, wsr2-1, wsr3-1, wsr4-2, irx1-6* (resistance control) and *agb1-1* (susceptibility control) plants were infected with the necrotrophic leaf pathogen isolate *Plectosphaerella cucumerina* BMM (*Pc*BMM). The relative fungal biomass was determined 5 days post infection (dpi) by qPCR analysis of the *Pc*BMM β-tubulin gene. Bars represent mean values and error bars indicate SD (n = 4-6). Asterisks indicate statistically significant differences to the wild type according to Student‘s t test (*p < 0.05; **p < 0,01).

## Discussion

In plants a mechanism is existing, which monitors and maintains the functional integrity of the cell walls (Doblas et al., 2018; Wolf, 2017). This mechanism seems to exhibit similarities to the one described in *S. cerevisiae* and is capable of detecting CWD and initiating adaptive changes in cellular and cell wall metabolism to maintain the functional integrity of the wall (Hamann, 2015). Understanding of the molecular mechanisms underlying CWD perception and the signaling cascades involved in regulating the CWD response in plants is increasing (Engelsdorf et al., 2018; Feng et al., 2018). However, our knowledge of the genes and molecular processes bringing about changes in cell wall metabolism in response to CWD and their function during growth and development is very limited. Here, we have determined if we can identify genes mediating responses to CWD in seedlings and cell wall metabolism in adult plants by combining seedling-derived transcriptomics data with FTIR-based cell wall analysis of seedlings with T-DNA insertions in selected candidate genes. This approach identified 14 genes (out of 23 original candidates), whose functions are not well understood and which belong to different gene families (Table S1). Very little is known about the biological function of *At1g74440* beyond that it encodes an ER membrane protein. The gene has been implicated in biotic and abiotic stress responses mediated by Plant Natriuretic Peptides (PNPs) based on co-expression with *AtPNP-A* (Meier et al., 2008). *ATFIM2* encodes a protein belonging to the Fimbrin family and seems to modulates the organization of actin filaments (Zhang et al., 2016). *SQE2* encodes a squalene epoxidase converting squalene into oxidosqualene, which forms the precursor of all known angiosperm cyclic triterpenoids (Rasbery et al., 2007). Triterpenoids are required for production of membrane sterols and brassinosteroids. *PSAT1* encodes an amino transferase required for Serine biosynthesis taking place in the chloroplast (Wulfert & Krueger, 2018). Serine biosynthesis in turn is required during photorespiration, a prerequisite for carbohydrate metabolism and plant growth. While AGPs have been implicated in cell wall remodeling, very little information is available regarding the specific function of AGP7 in this context (Seifert, 2018). *AtBXL1* encodes an enzyme acting during vascular differentiation as a β-D-xylosidase while acting as an α-L-arabinofuranosidase during seed coat development (Arsovski et al., 2009). *PRX47* encodes a putative peroxidase, is apparently expressed in differentiating vascular tissue in seedling roots and stems and involved in lignification (Tokunaga et al., 2009). *UGT71C3* and *UGT76B1* encode UDP-glycosyltransferases (UGTs), which have been implicated in glycosylation of phytohormones and / or metabolites during the response to biotic and abiotic stress (Rehman et al., 2018). UGT76B1 in particular seems to glycosylate isoleucic acid, which is required for coordination of SA- and JA-based defence responses active during infection by pathogens like *Pseudomonas syringae* and *Alternaria brassicicola* (Maksym et al., 2018). *ADR1-L1* encodes a coiled-coil nucleotide-binding leucine-rich repeat protein and forms an important element of the effector-triggered immunity in plants (Bonardi et al., 2011; Dong et al., 2016). Reviewing the available knowledge provides further evidence that several of the genes (*PRX47, SQE2, ATBXL1, AGP7, PSAT1*) are probably required for processes relevant for cell wall or plasmamembrane metabolism. Intriguingly *UGT76B1, UGT71C3, ADR1-L1* have been implicated before in the responses to abiotic or biotic stress, which also involves plant cell walls (Bonardi et al., 2011; Maksym et al., 2018; Rehman et al., 2018). In our experimental conditions the seedlings are exposed to CWD but not biotic / abiotic stress. Thus raising the possibility that these genes are actually responding to cell wall-related events, which may also occur during biotic and abiotic. More importantly the results suggest that the approach pursued here enables us to identify amongst the many genes in the Arabidopsis genome those that contribute to the responses to CWD and regulation of relevant aspects of cell wall and membrane metabolism.

We characterized four candidate genes in more detail. These had been selected based on the FTIR phenotypes apparently caused by insertions in the candidate genes and the limited detailed knowledge regarding their biological functions. qRT-PCR-based expression analysis of the four genes in ISX-treated seedlings yielded results similar to the data from the transcriptomics experiment. Experiments with loss- and gain-of-function alleles of THE1 showed that ISX-induced changes in the transcript levels of *WSR1, 2, 3* and *4* are sensitive to an increase in the activity of THE1 (*the1-4*) while effects of reductions (*the1-1*) are less pronounced (Merz et al., 2017). These results are to be expected since complete loss of THE1 results in reduced responses to CWD but not complete losses, suggesting that the THE1-mediated CWI maintenance mechanism is either redundantly organized or other signaling mechanisms exist (Engelsdorf et al., 2018). However, the results support the notion that *WSR* gene expression is regulated by the THE1-mediated CWI maintenance mechanism and that *WSR* activity might be controlled on the transcriptional level.

Table 2 provides a global overview of the phenotypes observed for the insertion lines in the four genes. Reduction of *WSR1* and *WSR2* activity seemed to cause similarly pronounced FTIR phenotypes in an area where diagnostic signals for cellulose and pectins are normally found (Figure 1). For *wsr1-2*, we detected increased amounts of cellulose, rhamnose and xylose in stem-derived material while glucuronic acid was reduced in leaf material (Figures 5, 6). ISX-induced SA production in seedlings and resistance to *Pc*BMM in adult plants were reduced (Figures 3, 7). In *wsr2-1* plants, we also detected an increase in cellulose in stem-derived material while responses to CWD, *Pc*BMM susceptibility and non-cellulosic cell wall matrix composition were similar to the controls (Figures 3, 5, 6, 7). Reductions in *WSR3* and *4* expression seemed to result in FTIR phenotypes related to cellulose elements, hemicelluloses and pectins (Figure 1). In *wsr3-1* stem-derived cell wall material, we detected significant differences in the amounts of fucose, rhamnose, arabinose and galactose compared to controls (Figure 6). In *wsr4-2*, cellulose content was reduced both in stem- and leaf-derived cell wall material, while the amounts of glucose and mannose in stem-derived material were reduced and increased, respectively (Figure 2, 5, 6). Analysis of responses to CWD found reduced JA and lignin production in ISX-treated *wsr4-2* seedlings and no differences to wild type in *wsr3-1* seedlings (Figures 3, 4). The phenotype observed in *wsr2-1* in combination with the limited available protein information provides unfortunately no new insights regarding the function of WSR2 (De Abreu-Neto et al., 2013). The specific effects on neutral cell wall sugars in *wsr3-1* plants suggest that WSR3 could contribute to cell wall polysaccharide metabolism possibly by mediating transport between the Golgi (where non-cellulosic cell wall polysaccharides containing fucose, rhamnose, arabinose, galactose are synthesized) and the plasmamembrane (Temple et al., 2016). This would make sense bearing in mind that WSR3 / SFH19 belongs to the SEC14-protein family, whose members have been implicated in phosphoinositide production required for membrane homeostasis and signaling processes regulating cellular processes like vesicle transport (Gerth et al., 2017; de Campos & Schaaf, 2017). These results implicated *WSR2* and *WSR3* in cell wall metabolism but not in the representative responses to CWD examined here. Both *WSR1/DIR7* and *WSR4/PXC3* seem to be required for CWD responses on the seedling levels and for specific aspects of cell wall metabolism in adult plants, suggesting that they are both involved in CWD-induced signaling processes regulating changes in cell wall composition. *WSR1* seems only required for increased SA production in response to ISX-induced CWD, whereas *WSR4* is required for both JA and lignin production. The *wsr4-2* phenotypes were similar to those described for CWD responses in *mik2* seedlings where also only JA and lignin production differ from controls while SA amounts are similar (Engelsdorf et al., 2018; Van der Does et al., 2017). These results suggest that both RLKs are required for the same aspects of CWI maintenance. The THE1-dependent reduction in *WSR4* transcript levels in response to CWD suggest it could repress CWD-induced responses. This would be similar to the CWD response phenotypes of *FER*, where a FER KD leads to enhanced production of JA, SA and lignin (Engelsdorf et al., 2018). However, the *WSR4* loss of function phenotypes suggest the RLK is required for ISX-induced JA and lignin production. This implies the existence of additional regulatory elements interacting with WSR4 to give rise to the observed mutant phenotypes. Since the related RLK PXC1 has been implicated in vascular development a function for WSR4 in coordination of CWD perception with cell wall metabolism is conceivable (Wang et al., 2013).

**Table 2:** Overview of the phenotypes observed for mutant lines of the different candidate genes examined. Statistically significant differences compared to the wild type are indicated with blue (increased) or red (decreased) arrows.

|  |  | *wsr1* | *wsr2* | *wsr3* | *wsr4* |
| --- | --- | --- | --- | --- | --- |
| Cell wall damage response<br>(6 day-old seedlings) | JA | - | - | - | ↓ |
|  | SA | ↓ | - | - | - |
|  | Lignin | - | - | - | ↓ |
| Cell wall composition<br>(5 week-old plants) | Cellulose | stem ↑ | stem ↑ | - | leaf ↓<br>stem ↓ |
|  | Fucose | - | - | stem ↑ | - |
|  | Rhamnose | stem ↑ | - | stem ↑ | - |
|  | Arabinose | - | - | stem ↑ | - |
|  | Galactose | - | - | stem ↑ | - |
|  | Glucose | - | - | - | stem ↓ |
|  | Xylose | stem ↑ | - | - | - |
|  | Mannose | - | - | - | stem ↑ |
|  | Galacturonic acid | - | - | - | - |
|  | Glucuronic acid | leaf ↓ | - | - | - |
| *PcBMM* susceptibility<br>(18 day-old plants) |  | ↑ | - | - | - |

To summarize, seedlings with T-DNA insertions in 14 of the 23 candidate genes that were selected in this study exhibited FTIR phenotypes. Gene expression analysis showed that *WSR* gene expression is modulated in response to ISX-induced CWD, with the modulation apparently sensitive to changes in THE1 activity. This connected the genes identified to the THE1-dependent CWI maintenance mechanism, suggesting that our approach has identified new components mediating CWI maintenance in Arabidopsis. Follow up studies with KO or KD lines for the four candidate genes found cell wall phenotypes in adult plants for all four and effects on CWD responses for *WSR1* and *4*. These results also suggest strongly that a more detailed analysis of the remaining 10 candidate genes identified, will probably yield interesting novel insights into the mode of action of the CWI maintenance mechanism and cell wall metabolism in general.

## Acknowledgments

This work was supported through Gatsby AdHoc funds and a grant from the Peder Sather Center for Advanced Study to T.H. and Chris Somerville. L.K. was supported by a Ph.D. Fellowship from the Porter Institute at Imperial College and I.C. by a PhD fellowship provided by the Royal Thai government. L.D. and A.W were supported through postdoctoral fellowships provided by the Porter Institute at Imperial College. T.E. was supported through a EU Marie Curie Fellowship “SUGAROSMO-SIGNALLING” and a DFG postdoctoral fellowship (EN 1071/1-1). N.G. was supported through the EEA project grant CYTOWALL. Research by A.M. was supported by Spanish Ministry of Economy and Competitiveness (MINECO) grant BIO2015-64077-R. Programming support from Ane-Kjersti Vie and help with lignin and phytohormone quantification from Trude Johansen and the PROMEC facility at NTNU are gratefully acknowledged. The authors declare they have no conflicts of interest.

## Acknowledgments

Programming support from Ane-Kjersti Vie and help with lignin and phytohormone quantification from Trude Johansen and the PROMEC facility at NTNU are gratefully acknowledged.

