## Supplemental Information for "Functional characterization of genes mediating cell wall metabolism and responses to plant cell wall integrity impairment"

**Table S1:** 23 candidate genes selected for follow studies based on gene expression profiling (Hamann et al., 2009 and Genevestigator). Gene annotations are based on Araport11. T-DNA insertion lines used for FTIR spectroscopy-based screening and functional characterization are listed. Genes in bold were selected for follow up studies.

| AGI | Affymetrix Probe Set | Gene Annotation | T-DNA Insertion Lines |
|---|---|---|---|
| *At1g07260* | 256053_at | UDP-GLUCOSYL TRANSFERASE 71C3 | SALK_021979 |
| *At1g74440* | 260211_at | ER membrane protein, putative (DUF962) | SALK_059087 |
| *At2g02950* | 266745_at | PHYTOCHROME KINASE SUBSTRATE 1 | SALK_005340 |
| *At2g13790* | 264107_s_at | SOMATIC EMBRYOGENESIS RECEPTOR-LIKE KINASE 4, BAK1-LIKE 1 | SALK_057955 |
| *At2g20010* | 265583_at | Gls protein (DUF810) | SALK_004645 |
| ***At2g35730*** | 265796_at | Heavy metal transport / detoxification superfamily protein | SALK_058271 (*wsr2-1*)<br>SALK_123509 (*wsr2-2*) |
| **At2g41820** | 260494_at | PXY/TDR-CORRELATED 3, Leucine-rich repeat protein kinase family protein | SALK_082484 (*wsr4-1*)<br>SALK_121365 (*wsr4-2*) |
| *At3g09010* | 259213_at | Protein kinase superfamily protein | SALK_116262 |
| *At3g11340* | 256252_at | UDP-DEPENDENT GLYCOSYLTRANSFERASE 76B1 | SAIL_1171_A11 |
| ***At3g13650*** | 256781_at | Disease resistance-responsive (dirigent-like protein) family protein | SALK_046217 (*wsr1-1*)<br>SALK_092919 (*wsr1-2*) |
| *At3g16560* | 258437_at | Protein phosphatase 2C family protein | SALK_023206 |
| *At4g33300* | 253377_at | ADR1-LIKE 1 | SAIL_302_C06 |
| *At4g33420* | 253332_at | PEROXIDASE 47 | SM_3_37097 |
| *At4g35630* | 253162_at | PHOSPHOSERINE AMINOTRANSFERASE 1 | SALK_074264 |
| *At5g24140* | 249773_at | SQUALENE MONOOXYGENASE 2 | SALK_012094 |
| *At5g24430* | 249730_at | Calcium-dependent protein kinase (CDPK) family protein | SALK_028536 |
| *At5g40760* | 249372_at | GLUCOSE-6-PHOSPHATE DEHYDROGENASE 6 | SALK_016157 |
| ***At5g47730*** | 248769_at | SFH19, Sec14p-like phosphatidylinositol transfer family protein | SALK_039575 (*wsr3-1*)<br>SALK_079548 (*wsr3-2*) |
| *At5g48460* | 248656_at | ATFIM2 | SALK_019403 |
| *At5g49360* | 248622_at | BETA-XYLOSIDASE 1 | SALK_054483 |
| *At5g56540* | 247965_at | ARABINOGALACTAN PROTEIN 14 | SALK_096806 |
| *At5g60660* | 247586_at | PLASMA MEMBRANE INTRINSIC PROTEIN 2;4 | SM_3_20853 |
| *At5g65390* | 247189_at | ARABINOGALACTAN PROTEIN 7 | SALK_039285 |

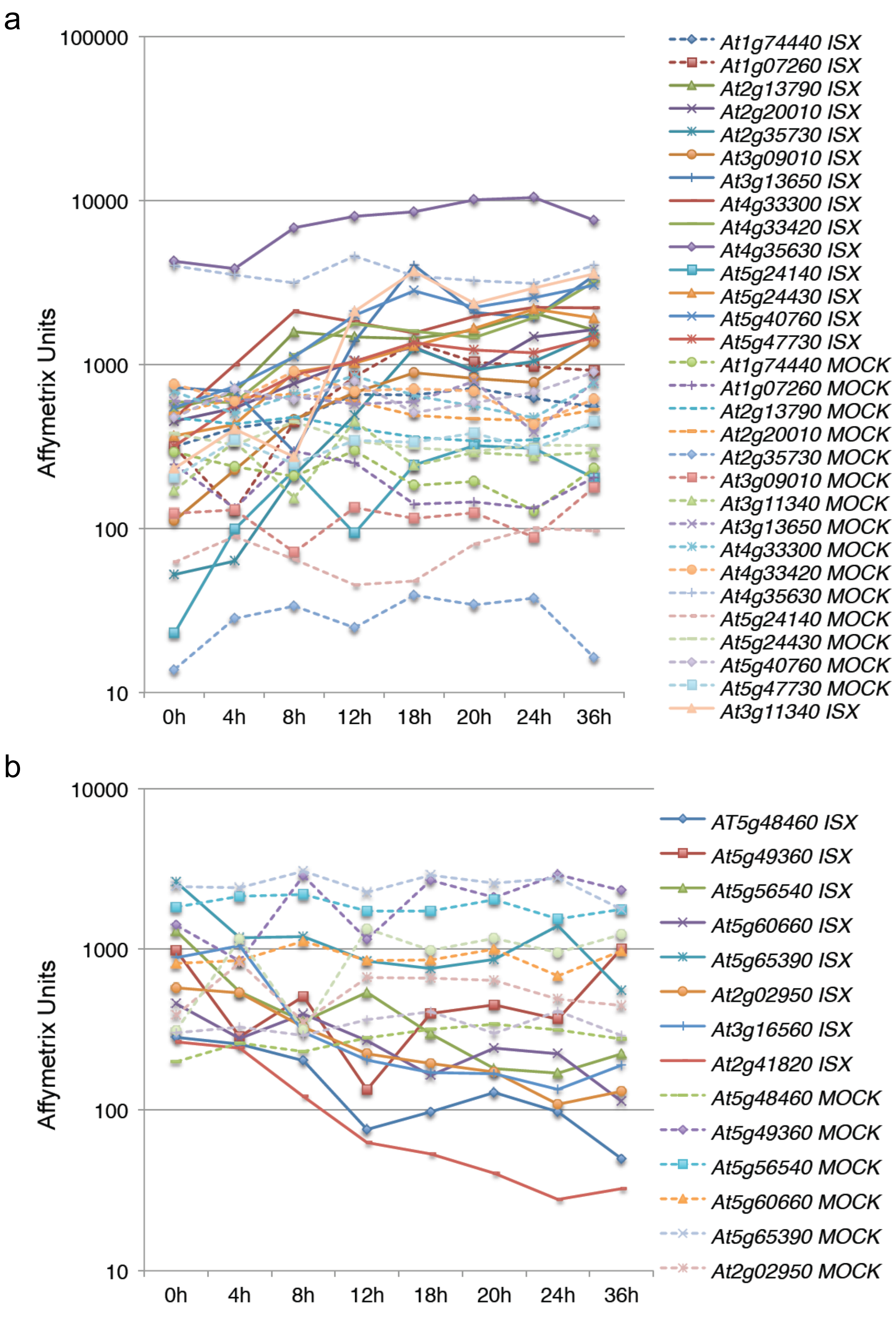


**Figure S1: DNA microarray-based expression analysis of 23 candidate genes.** Gene expression was monitored in 6 day-old Col-0 seedlings mock- (dashed line) or ISX-treated (solid line). **(A)** Genes where transcript levels are apparently elevated by isoxaben treatment. **(B)** Genes where transcript levels are apparently reduced by isoxaben treatment. Y-axis indicates transcript levels in affymetrix units, X-axis indicates time after treatment in hours.

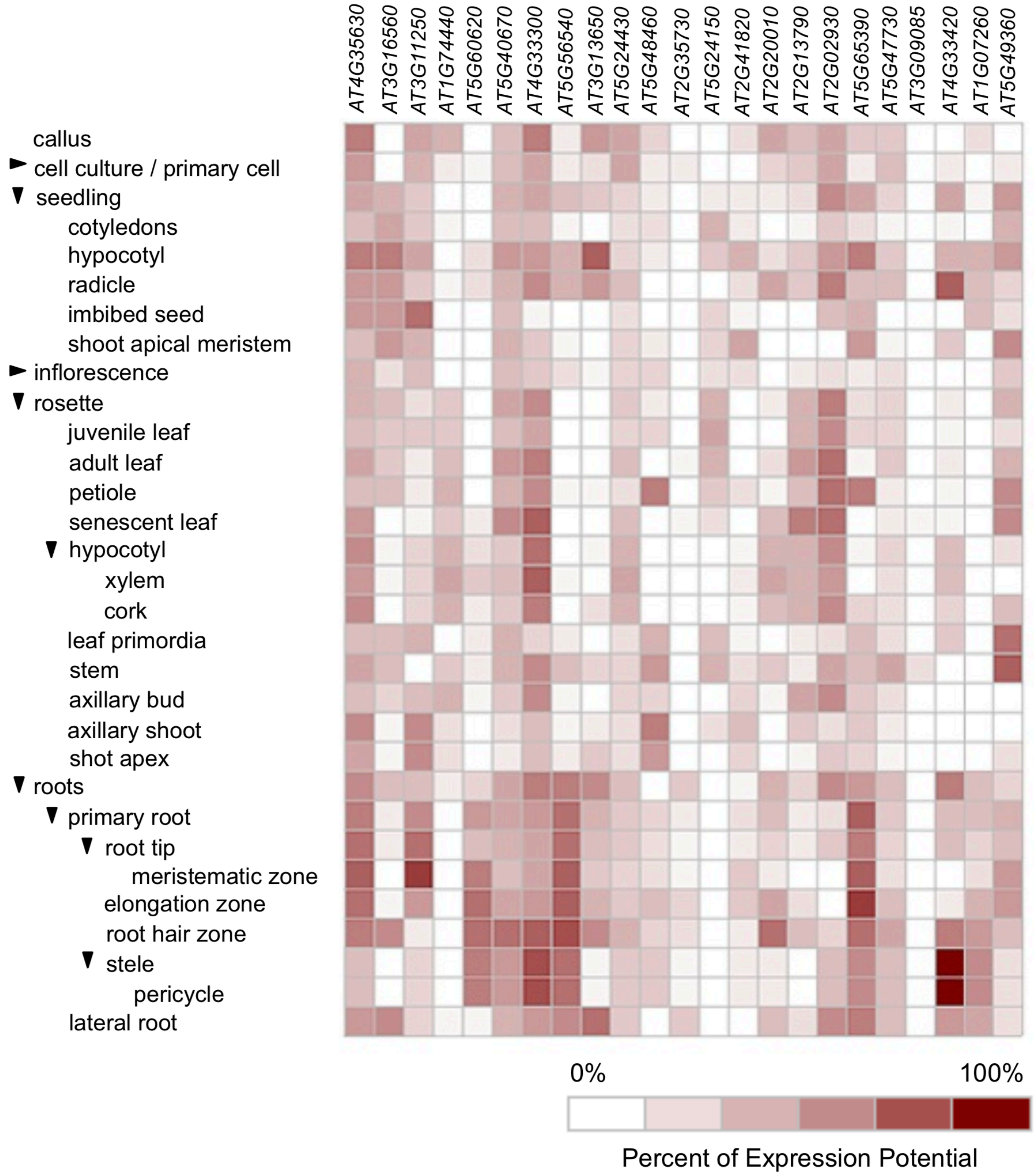


**Figure S2:** Graphic representation of the transcript levels for 23 candidate genes in different cell types and plant organs of *Arabidopsis thaliana* based on data deposited in the genevestigator database.

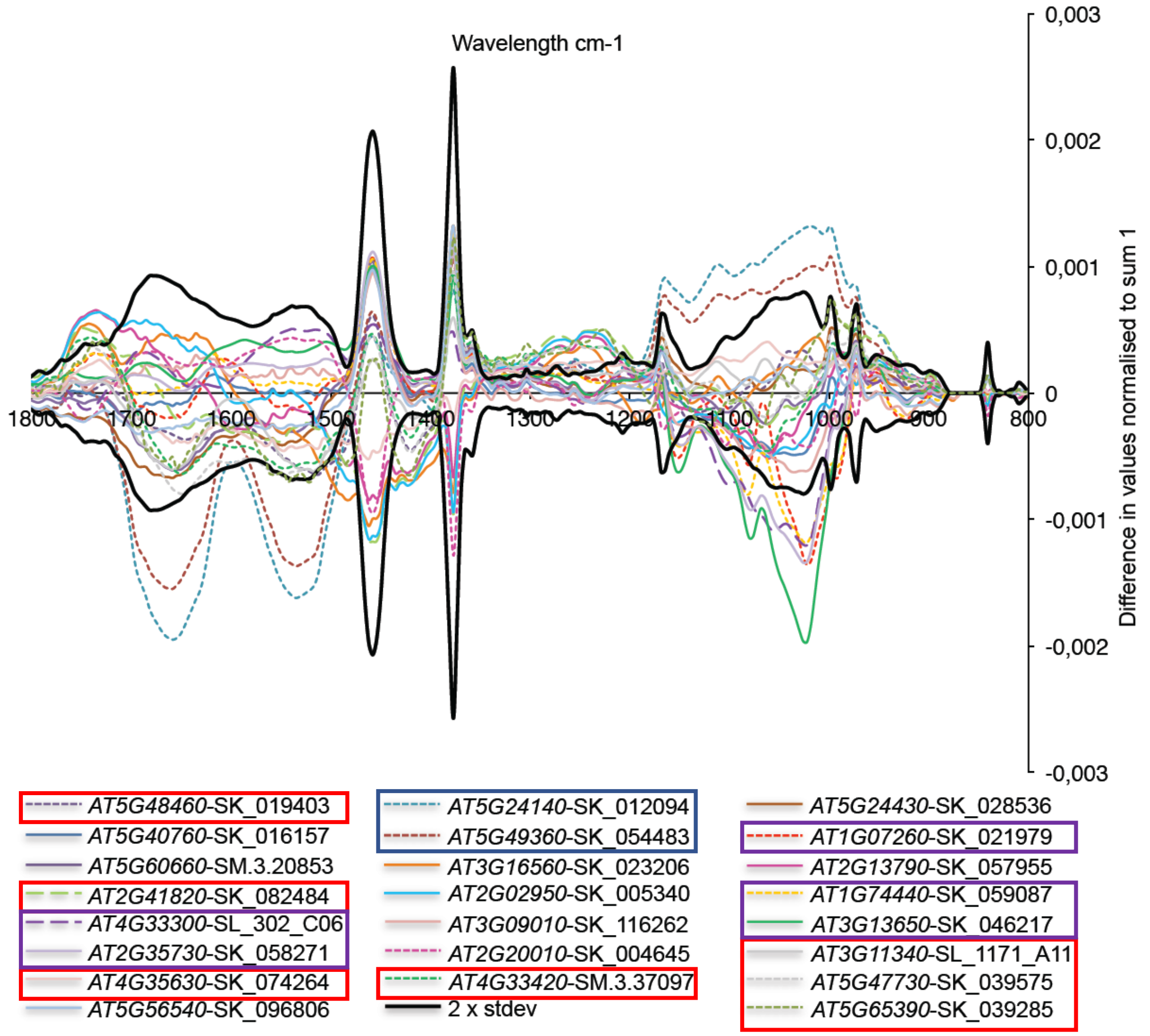


**Figure S3**. Graphic representation of average Fourier-Transform Infrared (FTIR) spectra of wild-type (Col-0) and all insertion lines screened. The black lines indicate 2x SD of the Col-0-derived seedling material. The different colored lines indicate data derived from the 23 insertion lines analysed. Colored boxes highlight genes where insertion lines exhibit similar FTIR-phenotypes.

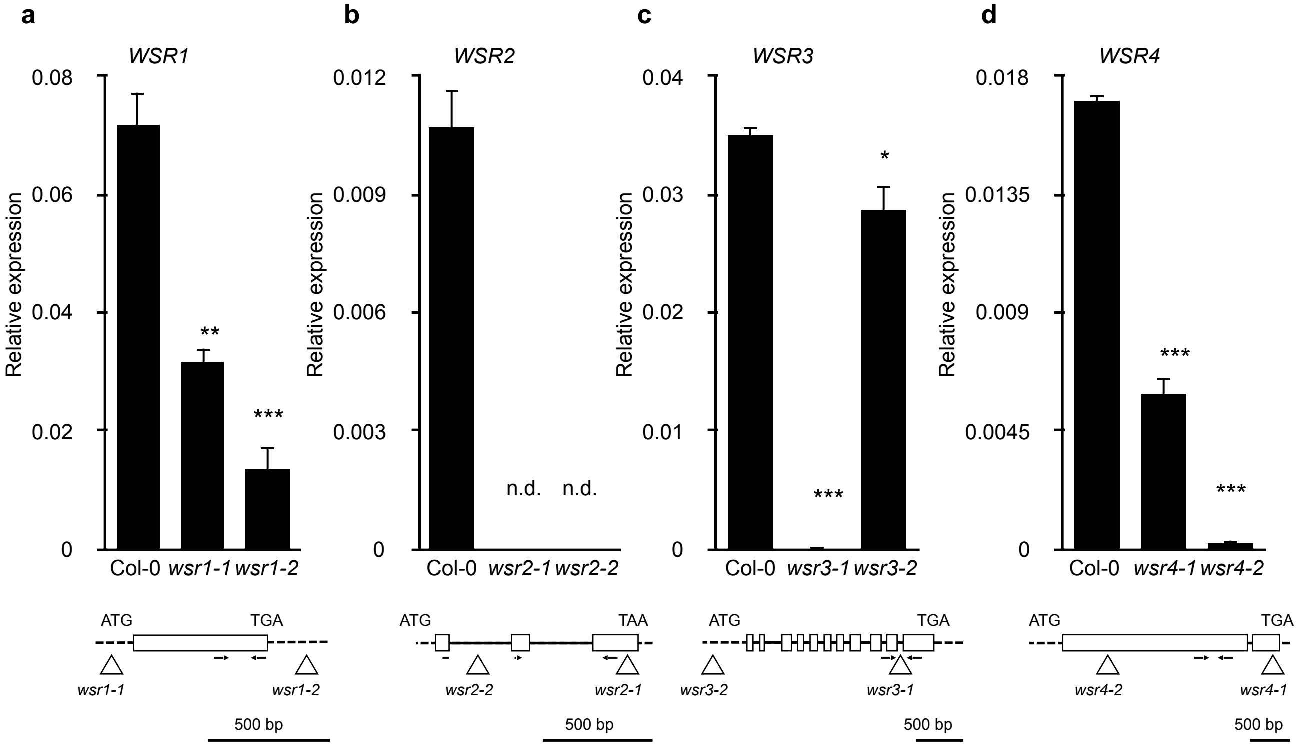


**Figure S4: Characterization of *WSR* T-DNA insertion lines.** Transcript levels of **(a)** *WSR1*, **(b)** *WSR2*, **(c)** *WSR3* and **(d)** *WSR4* were determined by qRT-PCR in the indicated *wsr* mutant seedlings. Values were normalized to *ACT2* and represent means from 3 independent experiments (n.d.: not detectable). Error bars indicate SD, asterisks indicate statistically significant differences to the wild type according to Student‘s t test (*p < 0.05; **p < 0,01; ***p < 0.001). Gene models indicating T-DNA insertion sites (triangles) and PCR primers used (arrows) are shown below the bar charts. Exons are represented by boxes, introns by lines and untranslated regions by dashed lines. Size bars indicate 500 bp.

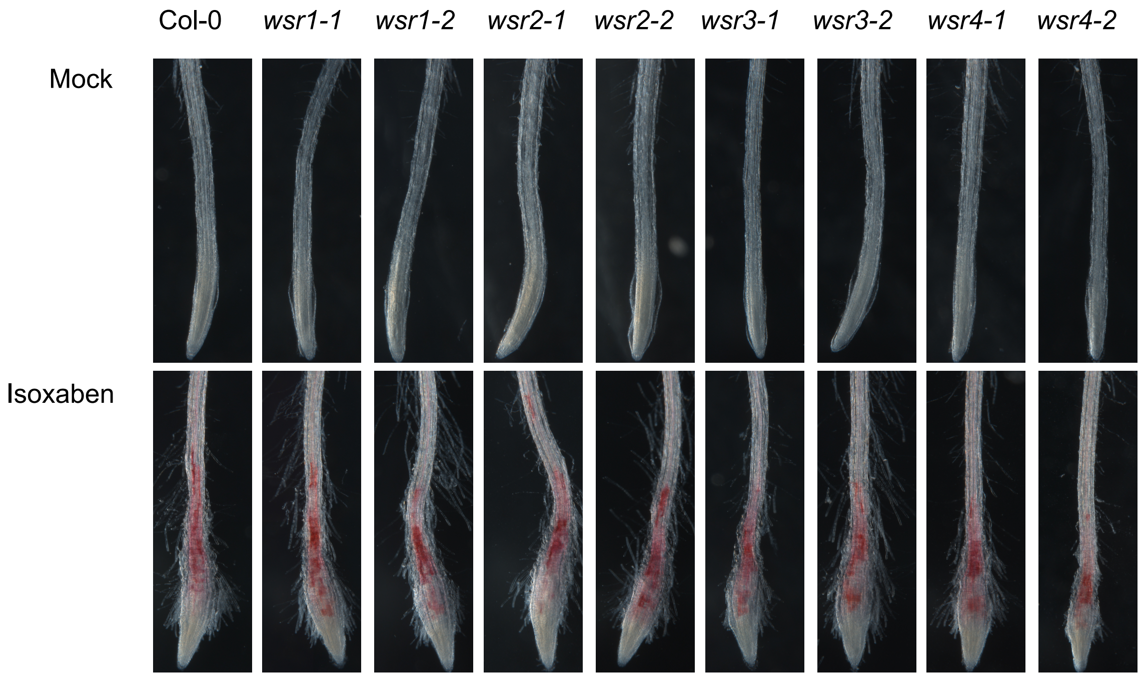


**Figure S5: ISX-induced lignification in root tips of candidate mutants.** Col-0, *wsr1-1*, *wsr1-2*, *wsr2-1*, *wsr2-2*, *wsr3-1, wsr3-2*, *wsr4-1* and *wsr4-2* seedlings were mock or ISX-treated for 24 h. Representitive images of phloroglucol-stained roots are shown.

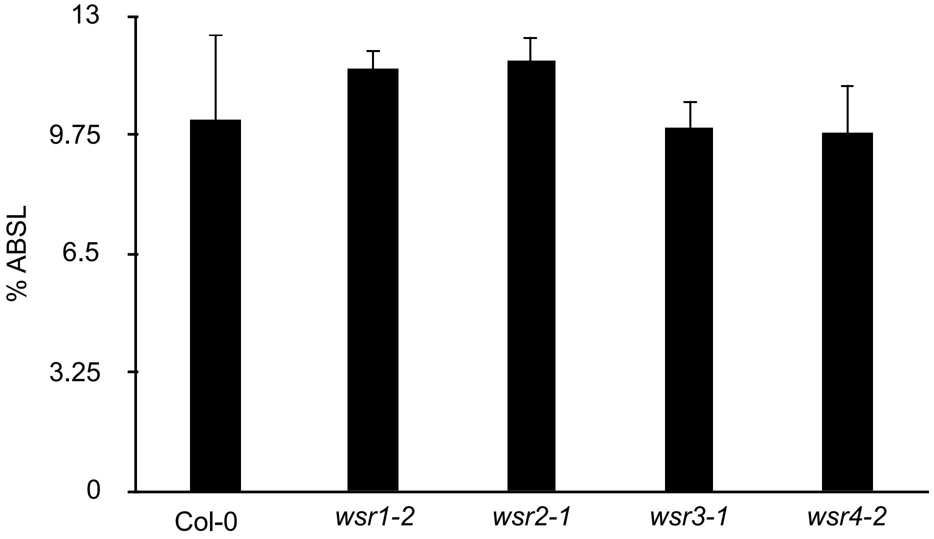


**Figure S6: Lignin content in stems of adult *wsr* plants.** Acetyl bromide soluble lignin was determined in stem cell wall preparations of 5 weeks-old Col-0, *wsr1-2*, *wsr2-1*, *wsr3-1* and *wsr4-2* plants. Bars represent mean values while error bars indicate SD (n= 4).
